# Achieving quantitative and accurate measurement of the human gut microbiome

**DOI:** 10.1101/2022.09.28.509972

**Authors:** Dylan Maghini, Mai Dvorak, Alex Dahlen, Morgan Roos, Scott Kuersten, Ami S. Bhatt

## Abstract

Robust benchmarking studies have highlighted how measured relative microbial abundances can vary dramatically depending on how DNA is extracted, made into libraries, sequenced, and analyzed. To build upon prior research, we investigated how sample preservation and storage choices impact observed absolute microbial load and relative metagenomic and metatranscriptomic measurements. Specifically, we studied how two common stool preservatives (OMNIgene GUT OMR200 and Zymo DNA/RNA PowerShield) perform across a range of storage temperatures (−80°C, 23°C and 40°C). For immediately frozen samples with no preservatives, we observed a mean colonic load of ∼100 trillion (1.2 × 10^14^) prokaryotes across ten donors, revising the gut prokaryote:human cell ratio of ∼1:1 to ∼4:1. We found that both preservatives introduce significant bias in the metagenomics results; and, while OMNIgene results were robust to storage temperature, samples stored in Zymo preservative had further bias with increasing storage temperatures. In terms of measured composition, we observed a ∼1.9x and ∼1.5x difference in the metagenomic Bacteroidetes:Firmicutes ratio in OMNIgene and Zymo preservatives, respectively. Absolute abundance measurements revealed that these differences are driven by higher measured Bacteroidetes in OMNIgene-preserved samples and lower measured Firmicutes in Zymo-preserved samples. For metatranscriptomic measurements, we also found that both preservatives introduced bias, but that RNA likely degraded in samples stored in OMNIgene preservative at high temperature. In summary, we recommend the OMNIgene preservative for studies that include significant field components. For metatranscriptomics studies, we recommend kits rated for RNA preservation such as the Zymo kit; however, existing samples collected in non-RNA rated kits might also be viable for limited metatranscriptomic studies. This study demonstrates how sample collection and storage choices can affect measured microbiome research outcomes, makes additional concrete suggestions for sample handling best practices, and demonstrates the importance of including absolute abundance measurements in microbiome studies.

## Introduction

Microbiome research relies on sequencing DNA or RNA to determine the relative abundances of various organisms, genes, or RNAs within a sample. It is known that different sample preservatives, storage conditions, DNA extraction methods, sequencing library preparation methods and bioinformatic analysis can impact measured relative abundances of microbes and microbial genes within a sample. However, the majority of studies, such as the robust and comprehensive Microbiome Quality Control project^1^, have only studied how DNA extraction, library preparation, sequencing and bioinformatic analysis choices impact results and overlook the impact of preservative choices and storage conditions. Relatively few, more limited studies, which are summarized in Supplementary Table 1, have reported how choice of sample preservative and storage temperature conditions can affect results. The results from these studies, while interesting, are at times conflicting, which makes it difficult to systematically determine the impact of preservatives and storage on measured microbial composition of a sample. Here, we evaluate the impacts of sample preservatives and handling on observed absolute microbial load and relative metagenomic and metatranscriptomic measurements.

User-friendly collection kits have gained popularity but produce similarly variable measurements as research grade preservatives. With the advent of home-collection kits such as OMNIgene GUT OMR200 (OMNIgene) and Zymo Research DNA/RNA Shield (Zymo) that are marketed for long-term room temperature storage and user-friendly collection, sample collection and preservation has become more reliable. Despite these advances, there have been discordant reports about the efficacies of these preservatives. A handful of studies have found that the OMNIgene and Zymo preservatives typically outperform other preservatives in recapitulating microbiome composition of immediately frozen samples^2–7^, which represent the current field standard for stool sample collection. By contrast, other studies have identified that these kits lower recovered taxonomic diversity or change abundances of various taxa^8–10^. While these kits, especially the OMNIgene kit, are extensively used, these preservatives have not been extensively validated at temperatures beyond room temperature, and it remains unknown whether taxonomic variations are due to microbial blooms during storage, biased taxonomic lysis, or biased depletion of nucleic acids. These kits have also not been validated and compared for RNA stability over extended time and temperature ranges that are typical for studies that involve sample collection at a site remote to the primary research location. Given the prevalent use of these preservatives, clear and robust studies are needed to understand how preservative use can bias microbiome analyses, in measurement of both relative and absolute abundances.

While most microbiome studies focus on relative abundance measurements, there is emerging evidence that measurement of the total count of microbes in the gut, or “absolute abundance”, provides a richer source of information. The use of absolute abundance measurements have been demonstrated to correct false conclusions drawn from relative data. For example, one study revealed that certain microbial taxa that appear relatively depleted in one soil environment are actually more abundant in absolute count due to a higher overall microbial abundance^11^. Absolute abundance measurements have also revealed key biological insights. For example, one study showed a ten-fold variation in total load across healthy individuals and a significantly lower microbial load in individuals with Crohn’s disease, while identifying multiple conclusions drawn from relative microbial profiling that were not maintained at the absolute level^12^. More recently, investigators using spike-ins of exogenous microbial cells to enable absolute quantification of microbes identified direct, exploitative interactions between gut bacteria and fungi in a preterm infant cohort during community assembly^13^. Methods such as microscopy, 16S rRNA FISH, spike-ins, and 16S rRNA qPCR can be used to quantify absolute levels of prokaryotes in the gut^14,15^; however, none are routinely used. This is unfortunate, as absolute quantification of microbes can prevent drawing artifactual correlations of microbes to one another and to biological outcomes, and can greatly inform the conclusions that are drawn about the effects of various components of the microbiome on each other and the human host. Incorporation of absolute quantitation relies on accurate and reliable measurement, however, little is known about the effect of preservative choice and storage conditions on the sample and resulting absolute measurements. This is particularly relevant as researchers are increasingly studying the gut microbiome in remote settings where cold chain for sample preservation cannot be easily maintained and thus using preservatives is necessary. Understanding the ‘real life’ consequences of preservative choice and transport temperatures on the measured microbial compositions of these samples is thus of critical importance.

Ideally, microbiome measurements should reflect the true state of the composition, abundance, and function of the gut microbiota. Unfortunately, it is currently unknown how sample collection methodology affects absolute abundance measurements. Even relative metagenomic and metatranscriptomic measurements have not been robustly evaluated at scale in certain common shipping and storage conditions. In an attempt to better fulfill this objective, we investigated the impact of several ‘real world’ preservation conditions on microbial measurements of stool samples. We evaluate storage conditions across ten different donor samples by quantifying the variation in microbial relative abundances at the genomic and transcriptional levels, and absolute prokaryotic abundances at the genomic level in OMNIgene and Zymo collection kits. We find an average total colonic load of 1.2 × 10^14^ bacteria (95% CI 5.1 × 10^13^ - 2.8 × 10^14^), which is approximately 3.2x higher than a previous estimate. By exposing samples to a range of storage conditions, we find that the use of either preservative leads to an absolute metagenomic and relative metatranscriptomic enrichment of Bacteroidetes and a depletion of Firmicutes, and we find that the OMNIgene preservative is most effective at stabilizing metagenomic sample composition when exposed to higher temperatures. Altogether, we expect that these sample preservation biases may lead to confounded microbial community measurements, and make concrete recommendations for specific best practices for future study design.

## Results

### Sample Collection and Study Design

Ten healthy adult donors from California, USA provided a single stool sample (Figure 1) as a part of a Stanford Institutional Review Board-approved research study. To evaluate the impact of storage temperature and preservative choice on measured stool microbial load and microbial composition, each sample was aliquoted either with or without a preservative buffer (OMNIgene GUT OMR200 collection tubes (OMNIgene) or Zymo Research DNA/RNA Shield Fecal Collection buffer (Zymo)). Samples without preservative buffer were immediately frozen at -80°C; samples with a preservative were either directly frozen at -80°C, or kept at either 23°C or 40°C for 7 days prior to storage at -80°C. Each of these seven experimental conditions was replicated in triplicate, for a total of 21 samples per participant (Figure 1).

**Figure 1:**
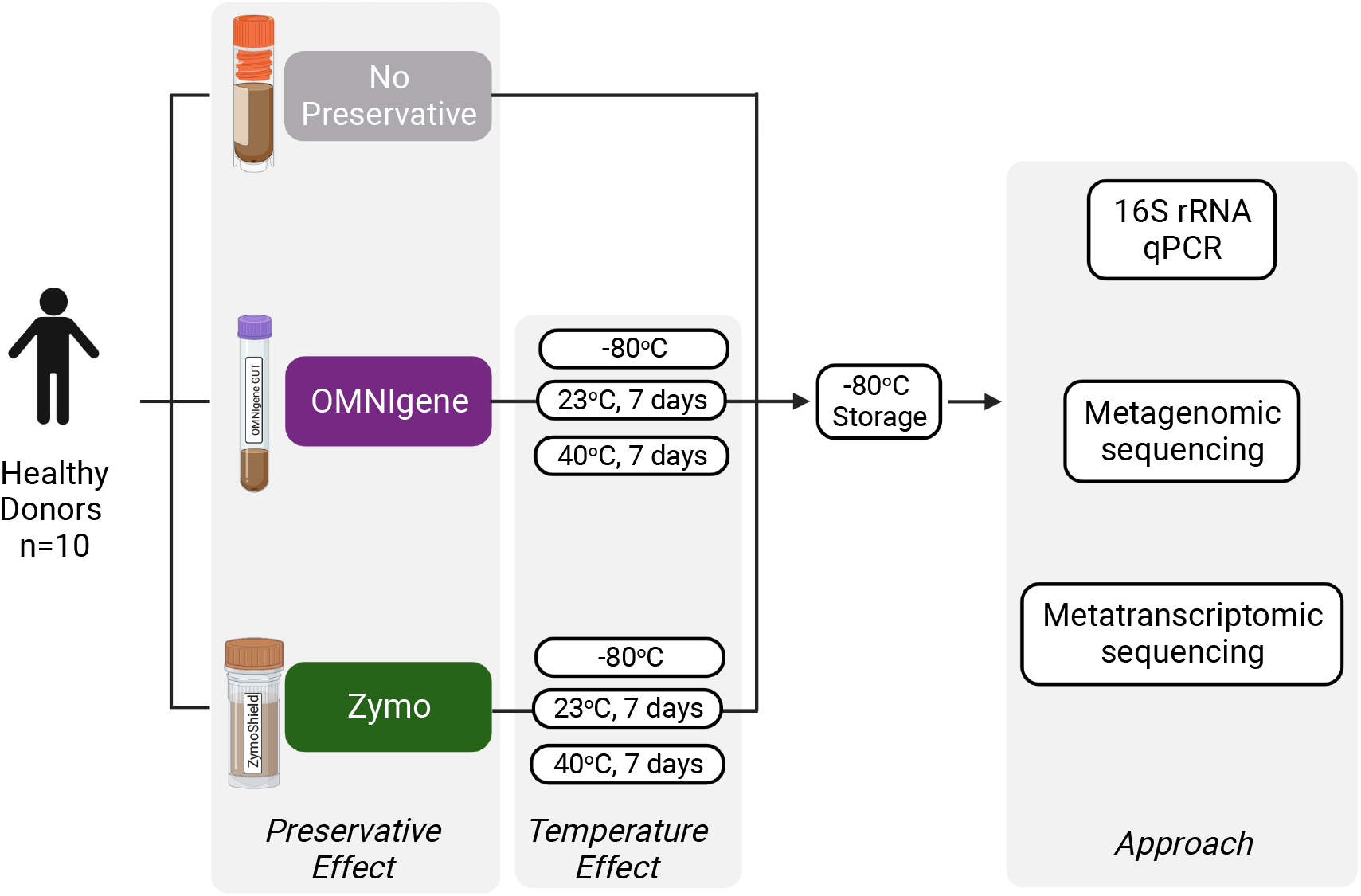
Overview of study workflow. Single stool samples were collected from ten donors. Each sample was stored in no preservative, DNA Genotek OMNIgene GUT OMR-200 preservative, or Zymo DNA/RNA Shield preservative. No preservative samples were stored immediately at -80°C. Samples in preservatives were stored at -80°C or stored for one week at 23°C or 40°C prior to storage at -80°C. All conditions were replicated in triplicate. Samples were then DNA extracted and RNA extracted, and measured with qPCR of the 16S ribosomal rRNA gene, metagenomic short-read shotgun sequencing, and metatranscriptomic short-read shotgun sequencing.

### DNA/RNA extraction, quality filtering and meta-’omic classification

DNA was extracted from 210 samples (Supplementary Figure 1) followed by 150 base pair (bp) paired-end sequencing, generating a median of 40.6 million reads per donor sample (range 11.3 - 231.5 million reads) (Supplementary Data 1-3) excluding one sample from Donor 3 stored in Zymo preservative at 40°C that failed library preparation. Median metagenomic read depth was 27.4 million reads (range 7.4 - 167.2 million reads) per sample after quality control (see Methods). RNA was also extracted from samples. While most samples yielded measurable RNA, we found that samples stored at 40°C in OMNIgene preservative, which is not rated for RNA preservation, did not reliably yield measurable RNA. Ribosomal RNAs were depleted from all samples. We performed 150 bp paired-end RNA sequencing on all samples that yielded RNA and generated a median of 62.1 million reads per donor sample (range 21.7 - 112.5 million reads) (Supplementary Data 4-6). Median metatranscriptomic read depth was 11.9 million reads (range 0.2 - 53.4 million reads) after quality control (see Methods). Quality-filtered metagenomic and metatranscriptomic reads were classified against a custom reference database encompassing microbial genomes in RefSeq and Genbank that were listed as “scaffold” quality or higher (as described in the methods). Classification results can be found in Supplementary Data 7 and Supplementary Data 8.

### Absolute Abundance of Gut Prokaryotes

We used qPCR targeting the bacterial/archaeal 16S ribosomal RNA gene to estimate the total prokaryotic load of the gut across conditions. Samples that were immediately frozen without preservative had an average of 1.33 × 10^12^ prokaryotes per gram of dry stool (95% CI 5.65 × 10^11^ - 3.13 × 10^12^) (Figure 2A; Supplementary Data 1-3). Adjusted for a previously reported total colonic volume of 400 mL^14^, this results in an estimate of ∼100 trillion (1.2 × 10^14^) total prokaryotes in a human gut, which is approximately 3.2x higher than a previous widely cited estimate^14^, although it is important to note that our estimate is based on ten donors. This estimate of total microbial load implies that the total prokaryote to human cell ratio is approximately 4:1.

**Figure 2:**
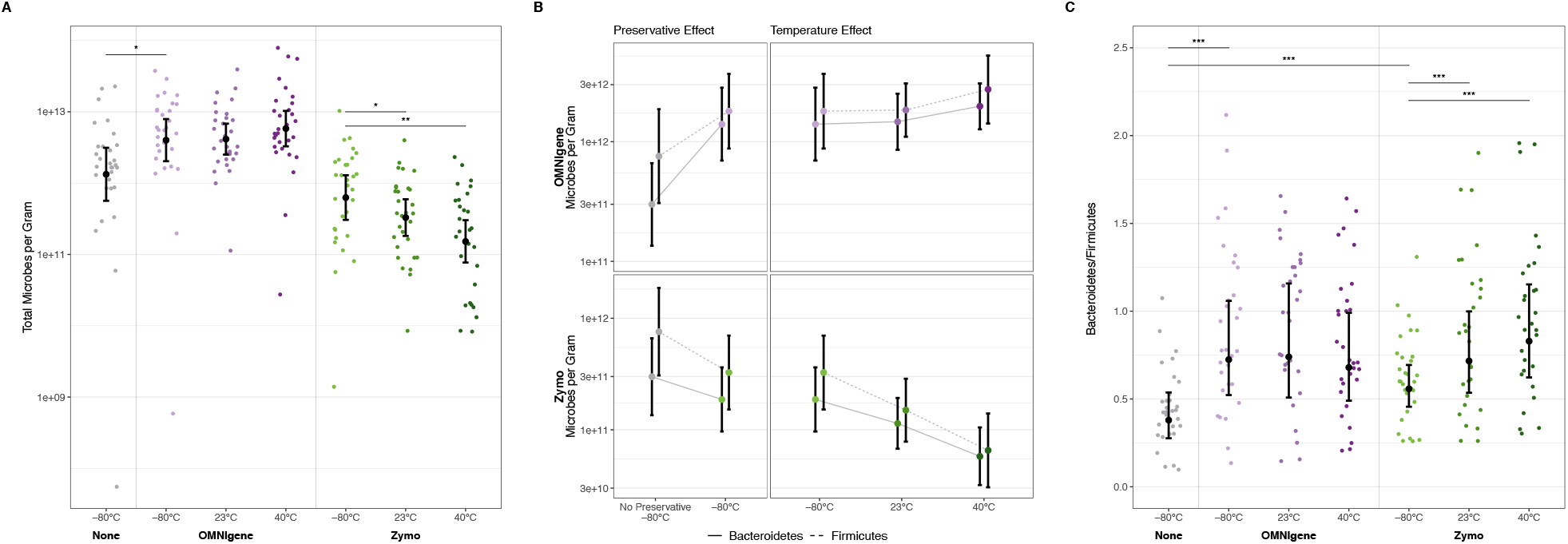
Absolute abundance quantification of microbiome samples. a) Total microbes per dry gram of stool for each sample as calculated with qPCR of the bacterial/archaeal 16S ribosomal rRNA gene. Scattered data points represent values from individual samples. b) Total count of Bacteroidetes and Firmicutes per dry gram of stool for each sample, in OMNIgene preserved samples (top) and Zymo preserved samples (bottom). Points represent estimated mean values from the GEE model. Significant differences are as follows: samples immediately frozen in OMNIgene preservative have an increase in Bacteroidetes relative to immediately frozen samples in no preservative (p=0.002). Samples stored at 23°C in Zymo have a decrease in Firmicutes (p=0.014) relative to samples stored in Zymo and immediately frozen. Samples stored at 40°C in Zymo have a decrease in Bacteroidetes (p=0.017) and Firmicutes (p=0.001) relative to samples stored in Zymo and immediately frozen. c) Ratio of Bacteroidetes count to Firmicutes count. Scattered data points represent values from individual samples. Center points indicate estimated mean values from the GEE model. Whiskers indicate 95% upper and lower confidence intervals from the GEE model. *p ≤ 0.05, **p ≤ 0.01, ***p ≤ 0.001.

These estimates of total microbial load were sensitive to how the samples were preserved and stored. Samples stored in OMNIgene preservative had a 200.9% higher observed microbial load relative to samples stored in no preservative (4.00 × 10^12^ bacteria per gram; 95% CI 2.03 × 10^12^ - 7.91 × 10^12^; p = 0.038), while samples stored in Zymo preservative had an insignificant but lower detected microbial load (6.27 × 10^11^ bacteria per gram; 95% CI 3.05 × 10^11^ - 1.29 × 10^12^; p = 0.141). Moreover, while samples in OMNIgene preservative had a similar measured bacterial load when stored either at 23°C or 40°C, samples stored in Zymo preservative yielded a progressively lower bacterial load when stored at higher temperatures (Zymo 40°C 1.52 × 10^11^ bacteria per gram; 95% CI 7.70 × 10^10^ - 3.02 × 10^11^; p = 0.004). Interestingly, preservative and temperature explain 37.6% of the variation in microbial abundance while donor explains only 25.9%, indicating that sample handling practices have greater influence than interindividual variation on absolute measurement. Together, these results suggest that OMNIgene buffer may lyse gut bacteria more effectively than standard extraction methods alone, while DNA may not be stable at higher temperatures when stored in Zymo preservative.

Using total estimates of absolute counts as well as metagenomic taxonomic abundance, we explored how storage preservative and temperature might differentially affect absolute counts of the three most abundant phyla in our dataset. We found that samples stored in OMNIgene preservative had no significant change in Actinobacteria load and higher total counts of Bacteroidetes and Firmicutes relative to immediately frozen, unpreserved samples (Supplementary Figure 2, Figure 2B), with a greater enrichment of Bacteroidetes relative to Firmicutes. Samples stored in Zymo preservative had a lower total load of Actinobacteria relative to immediately frozen, unpreserved samples (Supplementary Figure 2). With increasing temperature, samples preserved in OMNIgene showed similar degrees of enrichment across the three phyla, demonstrating the temperature stability of OMNIgene preservative. Conversely, samples stored in Zymo preservative had depletion of all three phyla with increasing temperature, with a greater depletion of Firmicutes relative to Bacteroidetes. Based on these changes in Firmicutes and Bacteroidetes absolute load with preservative and temperature, we considered the commonly reported ratio of Bacteroidetes and Firmicutes, which has been related to various health conditions (Figure 2C). We found that samples stored in OMNIgene preservative had a significantly higher Bacteroidetes:Firmicutes ratio (0.72; 95% CI 0.52 - 1.06; p ≤ 0.001) relative to unpreserved samples (0.38; 95% CI 0.28 - 0.54). Similarly, samples stored in Zymo preservative had a higher Bacteroidetes:Firmicutes ratio than unpreserved samples (0.56; 95% CI 0.46 - 0.69; p ≤ 0.001). Furthermore, the Bacteroidetes:Firmicutes ratio significantly increased as temperature increased in Zymo-preserved samples that were stored at 23°C or 40°C prior to freezing (0.72; 95% CI 0.54 - 1.00; p ≤ 0.001 for 23°C; 0.83; 95% CI 0.62 - 1.15; p ≤ 0.001 for 40°C). In summary, we observe that preservative choice has a strong effect on measured microbial load, emphasizing the impact of sample handling choices on absolute measurements. Further, we find that the use of absolute counts allows for the specific identification of which taxa contribute to changing relative ratios, demonstrating the importance of absolute abundance measurements in revealing key information that is otherwise obscured in relative abundance data.

### The Impacts of Temperature and Preservatives on Metagenomic Measurements

Storage temperature and preservative choice not only affect overall microbial abundance, but also affect the relative abundances of the most common microbes (Figure 3). We found considerable variation in relative community composition across the ten donors, but we also found systematic differences introduced by use of preservatives (Figure 3A). Furthermore, like our results for overall microbial abundance in the previous section, we found that samples stored with the Zymo preservative had additional systematic bias introduced when stored at higher temperatures. These results were consistent across a large number of different metagenomic measurements.

**Figure 3:**
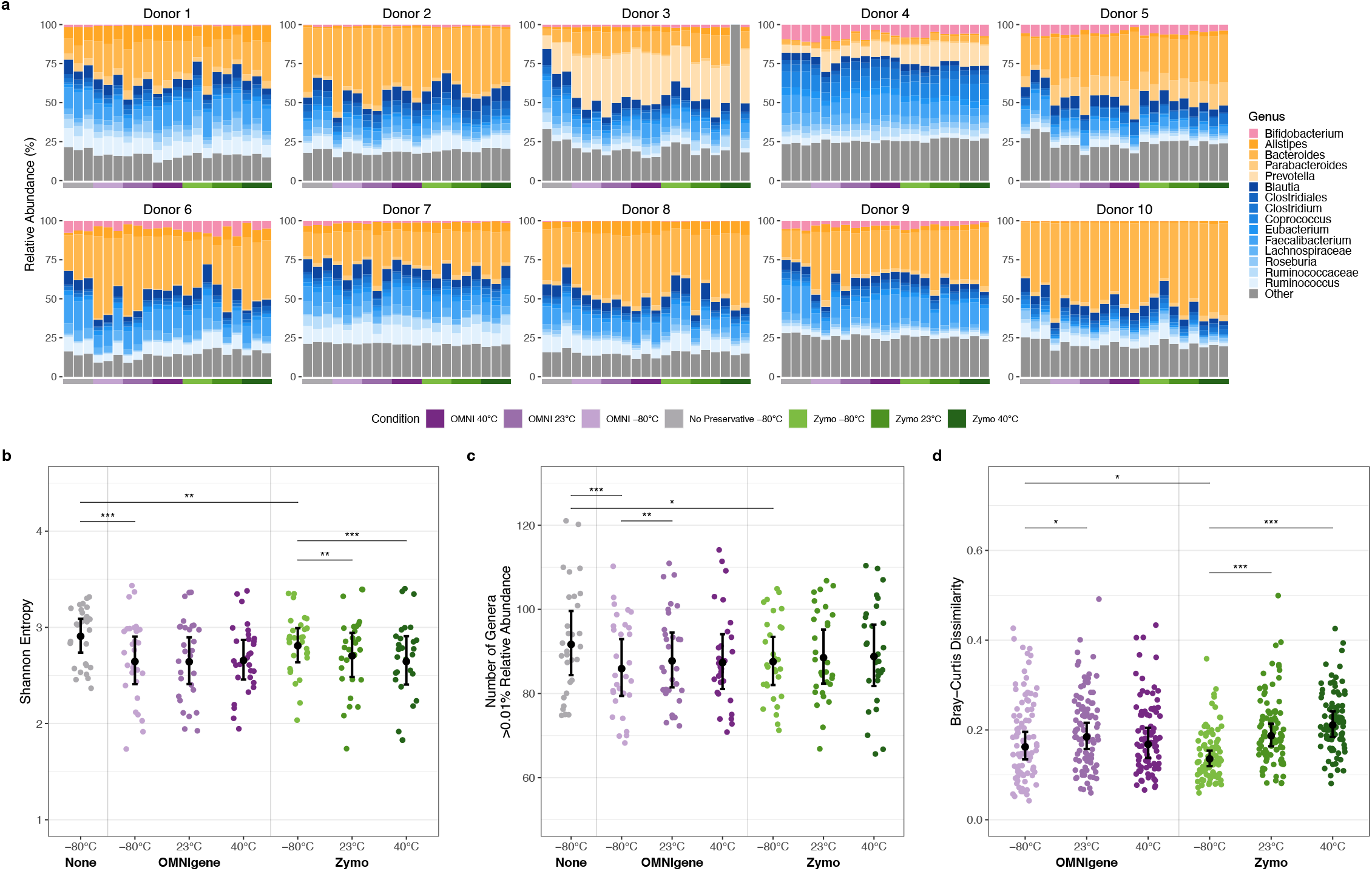
Metagenomic characterization of samples across storage conditions. a) Metagenomic relative abundance of the top 15 genera by total relative abundance across samples. Genera are colored by phylum. Only classified reads are shown. Replicate two from the Donor 3 Zymo 40C condition was excluded from the following analyses due to failed library preparation. b) Genus-level Shannon entropy across samples. Scattered data points represent values from individual samples. c) Geus-level richness across samples, filtered for genera present at greater than 0.01% v relative abundance. Scattered data points represent values from individual samples. d) Genus-level Bray-Curtis dissimilarity between samples from each preservative condition and the no preservative, immediately frozen condition. Each replicate from a given donor and condition was compared to each replicate from the corresponding donor in the no preservative, immediately frozen condition (for nine total comparisons per donor and condition). Scattered data points represent Bray-Curtis dissimilarity between pairs of samples. In panels B-D, center points indicate estimated mean values from the GEE model. Whiskers indicate 95% upper and lower confidence intervals from the GEE model. *p ≤ 0.05, **p ≤ 0.01, ***p ≤ 0.001.

Metrics of community diversity show significant differences across the preservation methods. Genus-level Shannon entropy was significantly lower in samples stored in either OMNIgene (2.7; 95% CI 2.4 - 2.9; p ≤ 0.001) or Zymo (2.8; 95% CI 2.6 - 3.0 p = 0.002) preservatives relative to unpreserved samples (2.9; 95% CI 2.7 - 3.1) (Figure 3B). As before, we found no significant changes with temperature for OMNIgene, but the measured entropy of samples stored in the Zymo kits progressively decreased when stored at 23°C (2.7; 95% CI 2.5 - 2.9; p = 0.009) and at 40°C (2.6; 95% CI 2.4 - 2.9; p ≤ 0.001). We observed similar trends using the inverse Simpson index (Supplementary Figure 3). We defined richness as the count of total genera present at >0.01% abundance. As with Shannon entropy, we found a decrease of -6.3% detected genera in OMNIgene-preserved samples (86 genera; 95% CI 79 - 93 p ≤ 0.001) and -4.5% detected genera Zymo-preserved samples (88 genera; 95% CI 82 - 93; p = 0.02) relative to unpreserved samples (92 genera; 95% CI 84 - 100) (Figure 3C). Richness was stable across temperatures in both preservatives, except for a slight increase in richness in samples stored in OMNIgene at 23°C relative to those that were immediately frozen (88 genera; 95% CI – 94; p = 0.007). Overall, we found that immediately freezing samples without a preservative is the best approach for maximizing detection of taxonomic diversity, as measured by Shannon entropy and overall richness.

To determine the similarity of the preserved samples to the immediately frozen, no preservative samples, we computed genus-level beta diversity (between sample differences) using the Bray-Curtis dissimilarity index formula (Supplementary Data 9-11). We found that samples stored in OMNIgene preservative were more dissimilar (median Bray Curtis dissimilarity of 0.16; 95% CI 0.13 - 0.20) than samples stored in Zymo preservative relative to immediately frozen, no preservative samples (median Bray Curtis dissimilarity of 0.14; 95% CI 0.12 - 0.15; p = 0.022) (Figure 3D). Further, we found that samples stored at 23°C and 40°C in Zymo preservative became increasingly dissimilar to immediately frozen, no preservative samples (0.19, 95% CI 0.16 - 0.21 for 23°C; 0.21, 95% CI 0.18 - 0.24 for 40°C; p ≤ 0.001 for both comparisons). Finally, we found minimal dissimilarity between technical replicates from the same sample, indicating that all storage methods had minimal technical variability (Supplementary Figure 4). In summary, we again find that both preservatives lead to shifts in community composition, and that increased temperature causes additional community shifts in the Zymo preservative.

Finally, we sought to examine how sample handling affects taxonomic relative abundances, as relative data are still commonly reported in the field. Specifically, we evaluated the relative abundances of the three most abundant bacterial phyla, Bacteroidetes, Firmicutes, and Actinobacteria, as well as viruses and fungi (Supplementary Figure 5). Compared to immediately frozen, no preservative samples, samples preserved in either OMNIgene or Zymo preservative showed significant relative enrichment of Bacteroidetes and a significant relative depletion of both Firmicutes and Actinobacteria. Also, while samples preserved in OMNIgene were robust to increasing storage temperature, we found that samples stored in Zymo preservative showed further enrichment of Bacteroidetes and depletion of Firmicutes and Actinobacteria as the storage temperature was increased. The only exception were viruses, which increased in abundance with temperature in both the OMNIgene and Zymo preservatives. We also tested for systematic biases introduced by preservative and storage temperature in all microbial genera with a relative abundance >0.1% in at least one condition (Supplementary Figure 6). Results seem to be driven predominantly by phylum-level effects: most Bacteroidetes genera, like Bacteroides and Alistipes, were enriched and most Firmicutes genera, like Faecalibacterium and Ruminococcus, were depleted. When stored at higher temperatures, Bacteroidetes genera were further enriched in the Zymo preservative, and Firmicutes genera showed heterogeneous responses to temperature in both preservatives. After adjusting for multiple comparisons, we found no statistically significant genus-level effects that were not already captured by the phylum-level effects characterized above.

Taken together, across a wide array of measured metrics related to taxonomic abundances, we see that both OMNIgene and Zymo preservatives lead to significant systematic differences from the immediately frozen, no preservative samples. Furthermore, while results in OMNIgene preservative are robust to temperature, Zymo kits show additional systematic differences when stored at higher temperatures. These results suggest that OMNIgene preservative differentially lyses some bacteria relative to unpreserved samples, while Zymo preservative better captures the composition of unpreserved samples but does not maintain sample composition with exposure to high temperature.

### The Impacts of Temperature and Preservatives on Metatranscriptomic Measurements

Metatranscriptomic analyses can quantify the active functional landscape of the gut microbiota, offering insight into the dynamic gene expression of gut microbes as they respond to environmental stimuli. While microbial transcriptional responses may be more compelling biomarkers of disease states, stabilization of RNA from stool samples is more difficult than stabilization of DNA because of the temperature sensitivity of RNA and the presence of potent RNases in stool samples. Unlike the Zymo kit, the OMNIgene kit is not rated for RNA preservation; however, it is among the most commonly used preservatives in stool microbiome studies, which means that scores of biobanked samples are preserved in this buffer. As there is likely interest in determining whether these samples may be extended for use beyond DNA-based applications, we evaluated the OMNIgene kit for its ability to preserve RNA for metatranscriptomic studies. We found that RNA could be extracted from samples stored in OMNIgene preservative and immediately frozen or exposed to 23°C for one week, indicating some potential for this kit to be used for transcriptomic analysis (Supplementary Figure 1). By contrast, we were unable to extract RNA from samples stored in OMNIgene preservative that had been exposed to 40°C for one week, and therefore excluded those samples from the following analyses.

We measured the metatranscriptome of samples across all ten donors and six conditions (Figure 4). We observed strong variability in metatranscriptomic taxonomic composition across preservation conditions and temperatures (Figure 4A), underscoring the importance of identifying an adequate stabilizer for RNA preservation. Similar to our observations in the absolute abundance and relative metagenomic data, we found that using OMNIgene or Zymo preservatives had significant effects on the measured outcomes. In contrast to the metagenomic results, we found that neither preservative was robust to temperature effects, though the Zymo preservative does yield RNA after exposure to 40°C. We observed these trends across many different comparisons of transcriptional composition.

**Figure 4:**
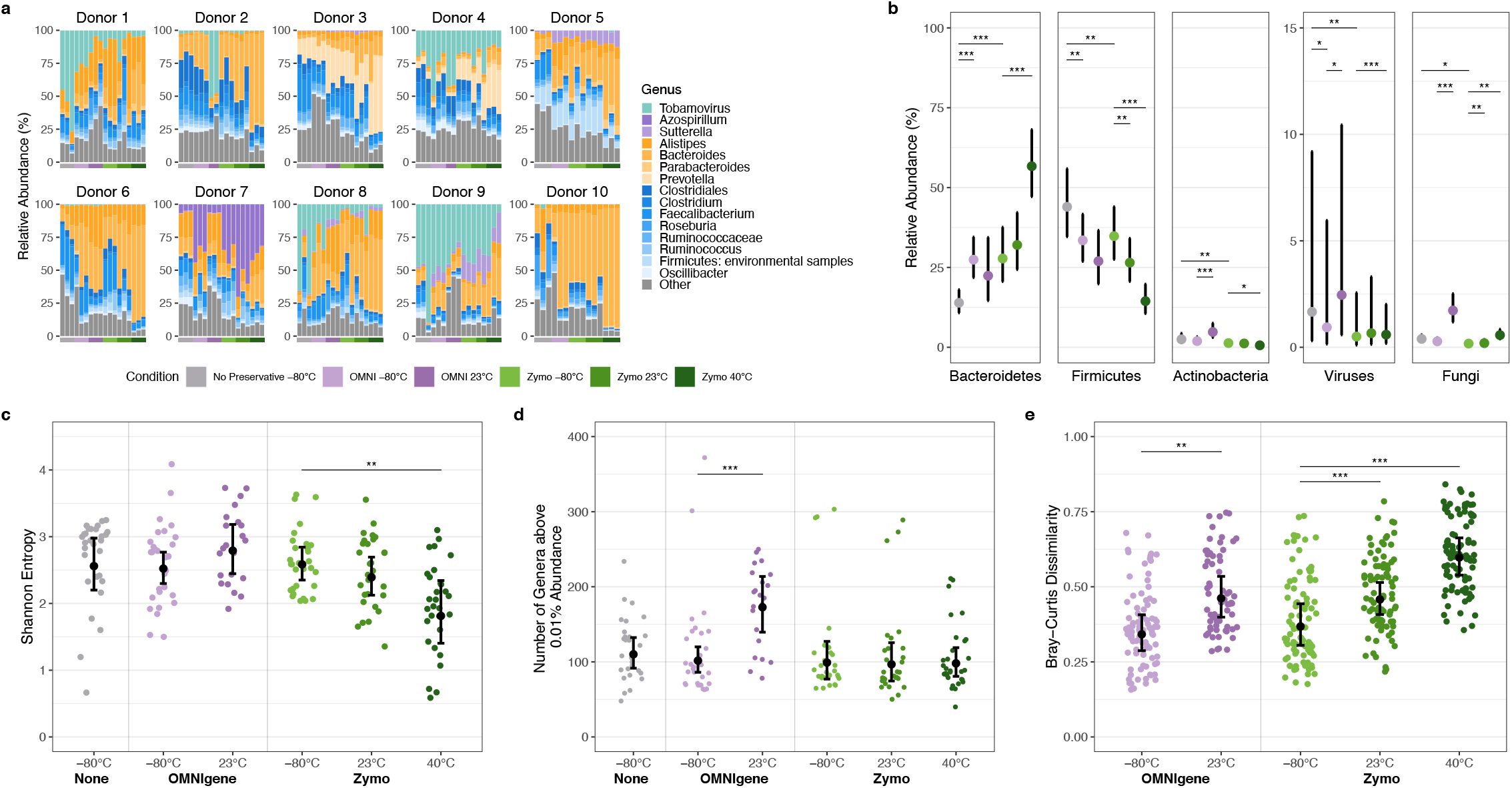
Metatranscriptomic characterization of samples across storage conditions. a) Metatranscriptomic relative abundance of the top 15 genera by total relative abundance across samples. Genera are colored by phylum. Only classified reads are shown. b) Metatranscriptomic relative abundance of the three most abundant bacterial phyla, viruses, and fungi across samples from each condition. c) Genus-level Shannon entropy across samples. Scattered data points represent values from individual sample. d) Geus-level richness across samples, filtered for genera present at greater than 0.01% relative abundance. Scattered data points represent values from individual samples. e) Genus-level Bray-Curtis dissimilarity between samples from each preservative condition and corresponding samples in the no preservative, immediately frozen condition. Scattered data points represent Bray-Curtis dissimilarity between pairs of samples. In panels B-E, center points indicate estimated mean values from the GEE model. Whiskers indicate 95% upper and lower confidence intervals from the GEE model. *p ≤ 0.05, **p ≤ 0.01, ***p ≤ 0.001.

First, we evaluated the relative abundances of transcripts from the most abundant bacterial phyla, viruses, and fungi to determine which specific microbial taxa were transcriptionally enriched or depleted across sample collection methods (Figure 4B). We found that samples immediately frozen in either preservative had a higher abundance of Bacteroidetes and a lower abundance of Firmicutes and viruses relative to immediately frozen, no preservative samples. Samples frozen in Zymo preservative had a lower abundance of Actinobacteria and fungi as well. Neither preservative was sufficient to protect against the effects of increased storage temperature, with all phyla demonstrating significant enrichment or depletion in at least one storage condition. We also tested for differences in metatranscriptomic relative abundance at the genus-level for all genera present at a relative abundance >0.1% in any condition. We observed that genus-level differences are largely driven by the phylum-level observations detailed above. Both OMNIgene and Zymo samples had a strong enrichment of Bacteroidetes genera such as Bacteroides, Parabacteroides, and Prevotella, and depletion of Firmicutes genera such as Faecalibacterium and Oscillibacter (Supplementary Figure 7). We also observed that immediately frozen Zymo samples had a strong depletion of the Tobamovirus virus, a RNA virus that infects tobacco, potatoes, tomatoes, and other crops.

Metrics of taxonomic community diversity showed more subtle differences across the preservation methods. Shannon entropy of samples stored in either preservative was comparable to no preservative, immediately frozen samples, and there were no significant changes with temperature in OMNIgene preserved samples (Figure 4C). By contrast, Shannon entropy was higher in the Zymo-preserved samples that were immediately frozen (2.59; 95% CI 2.35 - 2.84) compared to the Zymo-preserved samples that were exposed to 40°C (1.81; 95% CI 1.40 - 2.34; p = 0.003). Defining richness as the count of total genera present at >0.01% abundance, we found that samples stored in preservatives had similar richness relative to unpreserved samples (Figure 4D). Richness was stable across temperatures in samples stored in Zymo preservative, while richness increased by 10.7% in samples stored in OMNIgene at 23°C (173 genera; 95% CI 140 - 214) relative to those that were immediately frozen (101 genera; 95% CI 86 - 120; p ≤ 0.001). These metrics of alpha diversity demonstrate that samples stored in preservatives maintain similar alpha diversity to immediately frozen, no preservative samples, but preservatives have variable ability to protect against temperature changes.

We then compared beta diversity metrics to measure the microbial community similarity of preserved samples to immediately frozen, no preservative samples (Supplementary Data 12-14). Samples stored in either OMNIgene or Zymo preservative had increasing dissimilarity relative to the immediately frozen, no preservative samples when exposed to higher temperature (Figure 4E). Specifically, we observed higher dissimilarity in samples that were stored at 23°C in the OMNIgene (0.46; 95% CI 0.40 - 0.53) than those that were immediately frozen (0.34; 95% CI 0.29 - 0.41; p = 0.01), and observed higher dissimilarity in samples that were stored in Zymo preservative at 23°C (0.46; 95% CI 0.41 - 0.51) and 40°C (0.60; 95% CI 0.54 - 0.66) relative to samples that were stored in Zymo and immediately frozen (0.37; 95% CI 0.30 - 0.44; p ≤ 0.001 for both comparisons). Finally, we measured dissimilarity between technical replicates from the same sample and found that all storage methods had comparable technical variability (Supplementary Figure 4). Taken together, we observe that use of either the OMNIgene or the Zymo preservative leads to significant differences in measured metatranscriptomic composition relative to samples immediately frozen without preservative. We also observe that both kits, including those rated for RNA preservation, may still permit sample degradation that leads to significant shifts in metatranscriptomic taxonomic composition after exposure to high temperature.

## Discussion

Accurate measurement of the gut microbiome is essential for understanding the relationship between gut microbiota and human health. Such measurement relies on the investigation of a comprehensive range of variables and the minimization of study bias. Most studies measure relative taxonomic abundance, but overlook additional fundamental observations such as total microbial load and microbial transcript levels. Simultaneously, accurate measurement can be affected by biases in sample processing, from sample preservation through library preparation. Therefore, it is important to form data-driven decisions behind sample collection practices to optimize our ability to study the microbiome. Here, we focused on sample preservation, as variations in these practices can lead to nucleic acid degradation or biased microbial lysis^2,16,17^. We evaluate the performance of the two most common home collection kits for their ability to stabilize DNA and RNA and preserve total microbial load.

In comparing the performance of various preservatives, we observed differences in total microbial load, metagenomic alpha diversity and taxonomic relative abundances, and metatranscriptomic alpha diversity and taxonomic relative abundances. We posit that the majority of these differences occur as a result of two main phenomena: changes in lysis of microbial cells, and degradation of nucleic acids. Considering the detailed comparisons we report in this manuscript, we propose the following explanatory model: First, the addition of the OMNIgene preservative enhances overall lysis relative to unpreserved samples, and preferentially enhances lysis of certain organisms. This model is supported by the observation of a higher total microbial load in OMNIgene samples, and a further enrichment in total Bacteroidetes load relative to Firmicutes. We suspect that this increased load is due to improved lysis rather than degradation because we do not observe lowered total DNA concentration in samples exposed to higher temperatures and it is unlikely that no preservative, immediately frozen samples experience increased DNA degradation relative to immediately frozen OMNIgene samples, as established previously^6,18,19^. Consistent with the OMNIgene kit not being rated for RNA preservation, we find that the OMNIgene preservative does not protect against non-specific RNA degradation, as evidenced by lower or no RNA yield from samples exposed to higher temperatures. We observe a similar enrichment of Bacteroidetes and depletion of Firmicutes at the metatranscriptomic level as at the metagenomic level, reflecting the OMNIgene kit’s biased lysis of Bacteroidetes. Second, our model is that the Zymo preservative leads to biased lysis of certain organisms but does not protect as effectively against DNA degradation. This model is supported by the increased relative ratio of Bacteroidetes to Firmicutes, while diminishing total microbial load with increasing heat suggests that the Zymo preservative is not sufficient to protect against DNA degradation. We suspect that this change is due to nucleic acid degradation rather than decreases in lysis, as exposure to heat should not impair lysis. We also observe a relative enrichment of Bacteroidetes and depletion of Firmicutes taxonomic at the metatranscriptomic level, reflecting the Zymo kit’s biased lysis of Bacteroidetes.

Together, our results suggest that sample storage practices can lead to significant differences in observed microbial measurements that should be taken into account when designing experiments. As our model indicates that there is minimal DNA degradation of samples stored in the OMNIgene preservative, use of the OMNIgene collection kit may be advisable to reduce confounding for large cohort studies in which samples may travel for long periods or be exposed to high temperatures. Furthermore, the OMNIgene kit also yielded the highest total microbial load estimates, and is therefore recommended for absolute quantification studies. As our model suggests minimal RNA degradation in the Zymo kit and our results show a high degree of similarity between the Zymo kit and immediately frozen, unpreserved samples, the Zymo collection kit may be preferred for samples that will be immediately frozen after collection or for studies that plan on evaluating the metatranscriptome. These recommendations are detailed in Supplementary Table 2. Of note, this study was carried out before the introduction of a new DNA/RNA preservation product from DNAGenotek. Given the large number of studies that have already been carried out with the original OMNIgene kit, we anticipate that information on its performance for DNA/RNA applications as reported here will be useful for researchers who plan to access the likely hundreds of thousands of samples that have already been preserved in the OMNIgene reagent.

These results also help us frame the reproducibility crisis in the microbiome field. While many studies over the past decades have identified microbial features that have strong associations with human health outcomes, these associations are often inconsistent or not observed in follow up studies. Examples of this include discordant conclusions regarding the utility of the Bacteroidetes:Firmicutes ratio as a biomarker of dysbiosis^20^, the effects of Prevotella on gut inflammation and insulin sensitivity^21^, and the patient responsiveness to immunotherapy treatment after fecal microbiota transplant^22,23^. These disparate results may be due biological factors, such as microbial strain variation or patient-to-patient variation. However, the results herein suggest that technical noise due to sample collection and other aspects of study design can have a strong effect on observed microbial measurements. Furthermore, most existing studies rely on relative quantification, whereas we have observed that absolute quantification is necessary to disentangle true changes in overall microbial load and the individual taxonomic abundance. Therefore, we advocate that absolute microbial measurements should become standard practice in future microbiome studies.

While this study endeavored to be thorough in terms of assessing the role of preservatives and temperatures across multiple donors and replicates, it has several limitations. Our study was limited in both scope and size, focusing on ten donors from the United States who share relatively similar diets and lifestyles. We chose to maximize the number of conditions studied and technical replication over maximizing donor number. Further research in this space would benefit from evaluating more diverse cohorts to understand the effects of sample collection methods on a wider range of microbes. Additionally, while qPCR provided an estimate of total bacterial load, this method detects intact 16S rRNA gene sequences, which includes both dead and actively replicating bacteria and archaea while missing non-prokaryotic gut taxa, such as viruses and microbial eukaryotes. While absolute quantification can help disentangle whether taxonomic shifts are due to an increase in one taxon or decrease in another, we cannot definitively identify whether these shifts are due to changes in lysis efficiency, nucleic acid degradation, or microbial blooms. These results are based exclusively on extracted DNA, and therefore should be considered a lower bound as they do not reflect microbes that evaded lysis. Finally, there exist other sample preservation methods that are designed for RNA preservation that were not considered in this study. We chose to evaluate the OMNIgene GUT and Zymo DNA/RNA Shield kits as they represent two commonly used at-home stool collection systems, though the OMNIgene kits are not rated for RNA preservation. Emerging kits that are rated for both DNA and RNA preservation, such as the upcoming OMNIgene GUT DNA/RNA collection kit, can be evaluated using the framework we present here. Finally, we chose to focus this study on metagenomic and metatranscriptomic measurements, as these are common areas of investigation and compatible with many at-home stool collection kits. Future studies should incorporate other microbiome measurements, such as metabolomics, which represent an exciting and growing area of interest for microbiome researchers.

Through this study, we identified that sample collection methods can have a strong effect on microbial community measurements. We demonstrate that the use of preservatives can significantly affect total microbial load and alter genus- and phylum-level taxonomic abundances. Even in this small cohort of individuals, we found that the total absolute abundance of prokaryotes varied across donors. Given the importance of bacterial load and related features such as membrane lipopolysaccharide dosage on host biology and the importance of open niches for microbial community assembly, we suspect that total microbial load may be an important biomarker for disease progression and treatment outcomes. We expect that future research will leverage this method to measure absolute abundance to better correlate the microbiome and health outcomes. We anticipate that these results and related studies will guide best practices around large cohort study design, inform the cross-cohort comparisons made in meta-analyses, and enable researchers to optimize sample collection methods for their specific research questions.

## Methods

### Fecal sample collection

Fecal samples were collected from ten healthy adults in California, USA. Human subjects research approval was obtained (Stanford IRB 42043; PI: Ami S. Bhatt) and informed consent was obtained from all participants. Fecal samples were processed in three technical replicates across a range of conditions. These conditions include immediate storage at -80°C in no preservative, OMNIgene GUT OMR200 collection kits, or Zymo Research DNA/RNA Shield tubes, as well as storage at -80°C after temporary storage for seven days at 23°C or 40°C in the OMNIgene or Zymo collection kits. All stool was homogenized and stored at -80°C after processing.

### DNA extraction

All DNA extractions were performed using the QIAamp PowerFecal Pro DNA Kit (Qiagen). Sample input consisted of 250 mg of samples stored without preservative or 250 uL of samples stored in OMNIgene or Zymo preservatives. For each technical replicate, samples were randomly distributed across four batches of DNA extraction, such that the same donors or same conditions were not pooled together. Every extraction batch contained one blank negative control (sterile nuclease-free water) and one positive control (ZymoBIOMICS Microbial Community Standard). DNA extractions were performed according to the manufacturer’s protocol with the exception of using the EZ-Vac Vacuum Manifold (Zymo Research) instead of centrifugation. DNA concentration was measured using a Qubit 3.0 fluorometer (Thermo Fisher Scientific) with the dsDNA High Sensitivity kit.

### RNA extraction

All RNA extractions were performed using the RNeasy PowerMicrobiome Kit (Qiagen). Sample input consisted of 250 mg of samples stored without preservative and 250 uL of samples stored in OMNIgene or Zymo preservatives. Samples were extracted in the same batch randomization format as the DNA extraction. Every extraction batch contained one blank negative control (sterile nuclease-free water) and one positive control (ZymoBIOMICS Microbial Community Standard). RNA extractions followed manufacturer’s protocol with the exception of using the EZ-Vac Vacuum Manifold (Zymo Research) instead of centrifugation. RNA concentration was measured using a Qubit 3.0 fluorometer (Thermo Fisher Scientific) with the RNA High Sensitivity kit. Of note, samples stored in OMNIgene GUT and incubated at 40°C for seven days consistently failed RNA extraction (RNA levels were undetectable by Qubit). These samples were excluded from downstream analysis.

### Metagenomic library preparation and sequencing

Samples were split into 96-well three plates for library preparation. Each plate consisted of extractions from a single technical replicate (70 samples and associated extraction controls) that were randomly distributed across the plate. Metagenomic sequencing libraries were prepared using the Illumina DNA Prep Kit (Illumina, Inc.). Libraries from each plate were pooled in equal concentration (barring positive and negative controls, which were pooled in lower concentrations) and sequenced on a NovaSeq 6000 (Illumina, Inc.) at 2×150 reads.

### Metatranscriptomic library preparation and sequencing

Initial RNA cleanup was performed with RNAClean XP beads (Beckman Coulter) according to the Illumina Stranded Total RNA Prep (Illumina, Inc.)., with one additional EtOH wash. Samples were split into three 96-well plates for library preparation. Each plate consisted of extractions from a single technical replicate (70 samples and associated extraction controls) that were randomly distributed across the plate. Ribosomal rRNAs were depleted and metatranscriptomic libraries were prepared with the Illumina Stranded Total RNA Prep, Ligation with Ribo-Zero Plus Microbiome kit (Illumina, Inc.). Libraries from each plate were pooled in equal concentration (barring positive and negative controls, which were pooled in lower concentrations). We performed 150 bp paired-end RNA sequencing on all samples that yielded RNA using a NovaSeq 6000 (Illumina, Inc.).

### Metagenomic and metatranscriptomic preprocessing and profiling

Metagenomic reads from the same sample and replicate were merged and reads were filtered to have a minimum read length of 60, a minimum quality of 30, and trimmed using TrimGalore v0.6.5. Reads were deduplicated using SuperDeduper v1.2.0 with default parameters, and reads that aligned to the human genome were removed using BWA v0.7.17. Metatranscriptomic reads were trimmed and filtered for host reads using the same methods and parameters as above, excluding deduplication. Ribosomal RNA reads were removed using sortmerna v4.3.4 against RFAM and SILVA ribosomal RNA databases. Metagenomic and metatranscriptomic reads were classified using Kraken v2.0.9 against a custom reference database including GenBank bacterial and archaeal genomes assembled to “scaffold” quality or higher as of January 2020.

### 16S rRNA qPCR

Quantification of absolute abundance for each sample was determined by qPCR for all samples. For the qPCR reaction, all sample DNA was diluted 1:1000 in sterile nuclease-free water. Standards were created using custom-synthesized plasmids containing a portion of the 16S rRNA gene from either *F. prausnitizii* or *B. vulgatus* (Supplementary Table 3). These organisms were chosen as they are the most abundant organisms across all samples. Standards were diluted from 1:10-1:10M in sterile nuclease-free water to produce the 10-log-fold standard curve. Universal 16S rRNA primers, 331F/797R primers, were used, as previously described^24^. For each technical replicate, samples were randomly distributed across a 96-well plate, such that the same donors or same conditions were not pooled together. Each plate included two negative controls, DNA extraction buffer and sterile nuclease-free water, in duplicate.

PCR conditions follow the protocol described by Jian et al^25^. All qPCR samples were run in triplicate using the QuantStudio 12K Flex (Applied Biosystems) with SsoAdvanced Universal SYBR® Green Supermix (Bio-Rad). qPCR analysis was performed using QuantStudio Design & Analysis 2.6.0 (Thermo Fisher Scientific). To calculate total microbial load, Cq values for each sample were converted to the number of 16S rRNA copies per microliter using the standard curve. 16S rRNA copies per dry gram of stool was calculated by adjusting copies/uL by the total dry weight of stool present in each preservative and the total input volume for DNA extraction. The copies/gram were then divided by the sum of the relative abundance of a given taxon multiplied by its 16S rRNA copy number, as noted in rrNDB^26^, to yield total microbes per dry gram of stool. For taxa without a known copy number, the average 16S rRNA copy number across all taxa observed, 4.6, was used.

### Statistical analysis and plotting

Our statistical protocol was pre-registered with the Open Science Foundation prior to the completion of data collection (https://osf.io/vj2fx). Our primary outcomes were the abundance of the three most common bacterial phyla, Bacteroidetes, Firmicutes, and Actinobacteria, as well as the abundances of viruses and fungi. Each of these taxonomic abundances was measured in three ways: as an absolute abundance (microbes per gram of dry stool), as a metagenomic relative abundance, and as a metatranscriptomic relative abundance. We also reported the abundance ratio between Bacteroidetes and Firmicutes to test for disproportionate depletions between those two phyla. For our secondary outcomes, we focused on genus-level sequencing results and normalized by total classified matches. We considered several measures of alpha- and beta-diversity: Shannon entropy, Inverse Simpson distance, richness of genera with a relative abundance greater than 0.01%, and Bray-Curtis distance; we considered each of these diversity metrics separately for the metagenomic and metatranscriptomic data. Finally, we looked at each microbial genus with a relative abundance greater than 0.1% in at least one condition.

For each of these outcomes, we tested for systematic differences by kit and by temperature with a Generalized Estimating Equations (GEE) approach: we used an unadjusted regression model with fixed effects for the 7 conditions, and an exchangeable correlation structure between the participant-level clusters to account for repeat measurements. We varied our distributional assumption based on the different outcomes. Specifically, for absolute abundances, we used a log-transformed linear model which brings the distribution into a roughly Gaussian shape. For relative abundances, we followed the approach of MaAsLin2^27^ and used a log-transformed linear model on TSS-normalized data, which they found to be a robust method for handling inherently compositional data. We controlled for multiple comparisons in our secondary outcomes of the relative abundances of individual genera with a Benjimini-Hochberg correction with a false discovery rate of 10%, again following the methodology of MaAsLin2. To model the ratio of Bacteroidetes to Firmicutes, we transformed the ratio to a probability and log transformed. Finally, to model richness, we used a Poisson count model, and to model Bray-Curtis distance, we focused only on within-patient distances and used an unadjusted model with fixed effects for each pair of conditions that we measured distance between. The GEE models were fit using the statsmodels package (v0.13) of Python.

For a sensitivity analysis, we repeated this analysis for our primary outcomes using a different model structure: we took a within-patient approach instead of a marginal approach, replacing the GEE with patient-level fixed effects. Results were robust to these modeling changes (Supplementary Table 4).

Plotting was performed using R v4.1.2 with packages tidyverse v1.3.1^28^, reshape2 v1.4.4^29^, ggsignif v0.6.3^30^, ggplot2 v3.3.5^31^, cowplot v1.1.1^32^, ggpubr v0.4.0, ggnewscale v0.4.7^33^, and paletteer v1.4.0^34^. Figure 1 was created using BioRender.

## Data Availability

All sequencing data generated for this study will be deposited on the NCBI Sequence Read Archive prior to publication. Source data for figures is available on GitHub at https://github.com/dgmaghini/Benchmarking.

## Code Availability

Workflow for metagenomic and metatranscriptomic preprocessing can be found at https://github.com/bhattlab/bhattlab_workflows. Workflow for metagenomic and metatranscriptomic taxonomic classification can be found at https://github.com/bhattlab/kraken2_classification. Analysis and plotting scripts can be found at https://github.com/dgmaghini/Benchmarking. Python code for fitting the GEE models can be found at https://github.com/alex-dahlen/Gut_Microbiome_Measurement_Bias.

## Acknowledgements

We thank the study participants for their participation in this study. We thank Scott Hazelhurst, Ovokeraye Oduaran, and Gary Schroth for their thoughtful recommendations for this project. We thank Erin Brooks, Meenakshi Chakraborty, Alvin Han, Aravind Natarajan, Ryan Park, Summer Vance, and Soumaya Zlitni for their technical assistance and support. This work was supported in part by NIH grant P30CA124435, which supports the Stanford Cancer Institute Genetics Bioinformatics Service Center. Computing costs were also supported, in part, by a NIH S10 Shared Instrumentation Grant 1S10OD02014101. This work was supported in part by NIH R01AI14862302 & R01AI14375702, a Stand Up 2 Cancer Grant, the Chan Zuckerberg Initiative, a Sloan Foundation Fellowship and the Allen Distinguished Investigator Award (to A.S.B.). We thank David Solow-Cordero and Sopheak Sim for assistance in using the Stanford Functional Genomics Facility and High-Throughput Bioscience Center, which is supported by the NIH Shared Instrumentation Grant S10RR019513, S10RR026338, S10OD025004, S10OD026899 and by an anonymous donation. D.M. is supported by the Stanford Graduate Fellowships in Science and Engineering program and the Stanford Gerald J. Lieberman Fellowship. M.D. is supported by NIH Cellular and Molecular Biology Training Program training grant T32GM007276.

## Author Information

A.S.B., S.K., D.M., M.D., and A.D. conceptualized the study. D.M. and M.D. designed the study, enrolled participants, collected samples, and performed extraction and qPCR on all samples. S.K. and M.R. performed ribosomal RNA depletion, library preparation, and sequencing on all samples. D.M., M.D., and A.D. carried out all analysis and generated figures. D.M., M.D., A.D., and A.S.B. wrote the manuscript. All authors read and approved the final manuscript.

## Ethics declarations

Competing Interests: S.K. and M.R. are employees of Illumina, Inc.

