## Supplementary Figures for "Achieving quantitative and accurate measurement of the human gut microbiome"

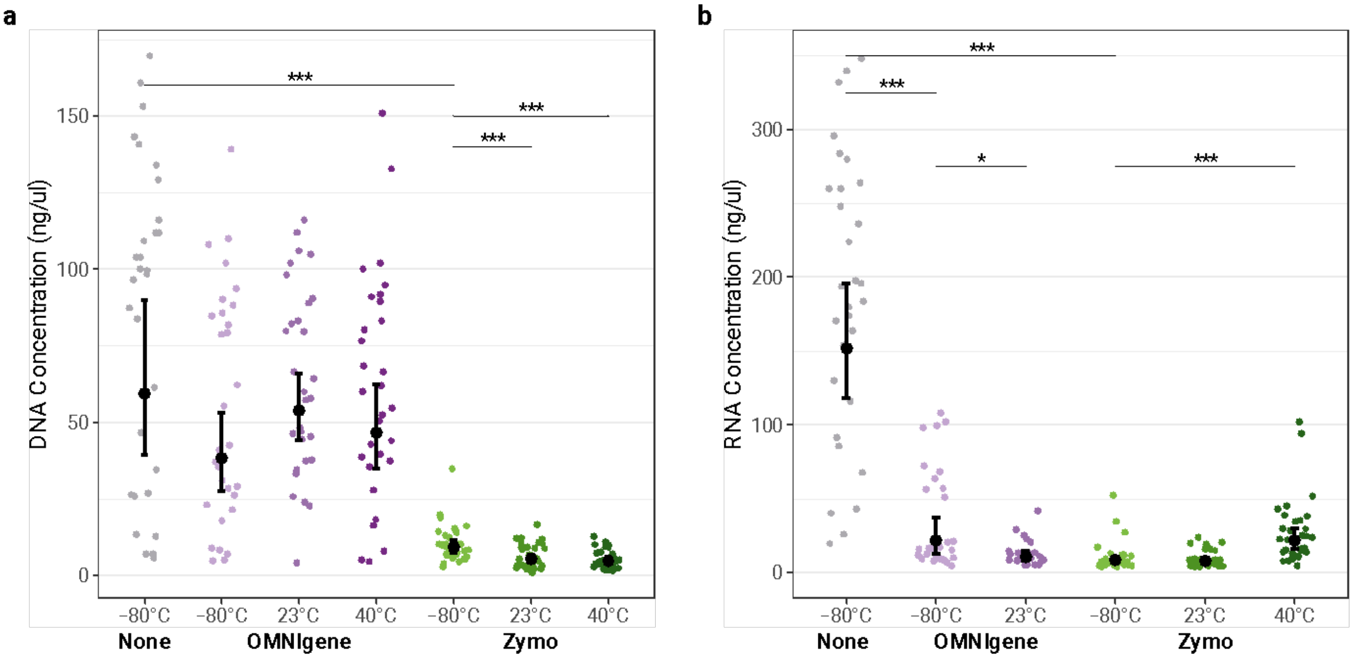


**Supplementary Figure 1: DNA and RNA extraction yields across conditions**

DNA concentration (ng / microliter) (a) and RNA concentration (ng / microliter) (b) of extractions across conditions. Samples that did not yield RNA are not shown. Scattered data points represent values from individual samples. Center points indicate estimated mean values from the GEE model. Whiskers indicate 95% upper and lower confidence intervals from the GEE model. *p ≤ 0.05, **p ≤ 0.01, ***p ≤ 0.001.

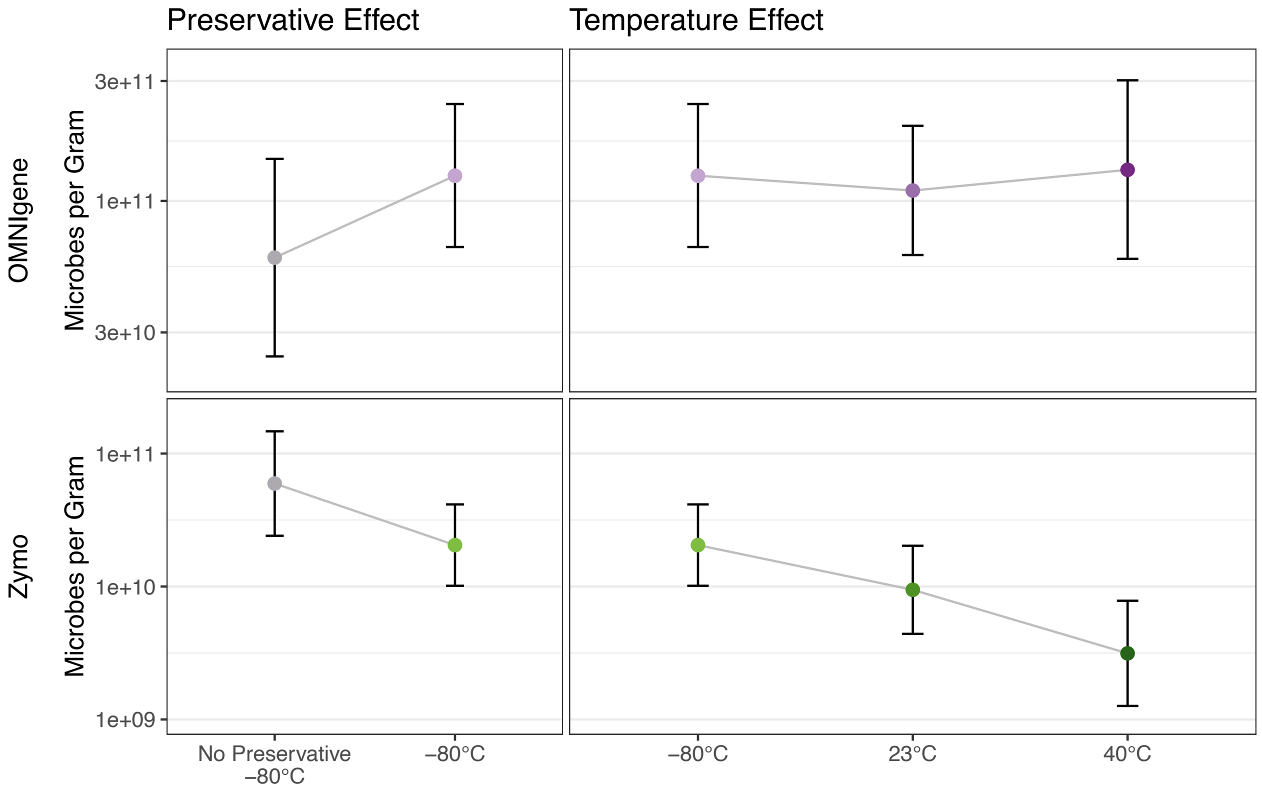


**Supplementary Figure 2: Absolute abundance of Actinobacteria**
Total count of Actinobacteria per gram of dry stool for each sample, in OMNIgene preserved samples (top) and Zymo preserved samples (bottom). Center points indicate estimated mean values from the GEE model. Whiskers indicate 95% upper and lower confidence intervals from the GEE model. Significant differences are as follows: samples immediately frozen in Zymo preservative have a decrease in Actinobacteria relative to immediately frozen samples in no preservative ($p=0.028$). Samples stored in Zymo preservative at 23°C and 40°C have a decrease in Actinobacteria relative to samples stored in Zymo preservative and immediately frozen ($p = 0.014$ and $p \leq 0.001$, respectively).

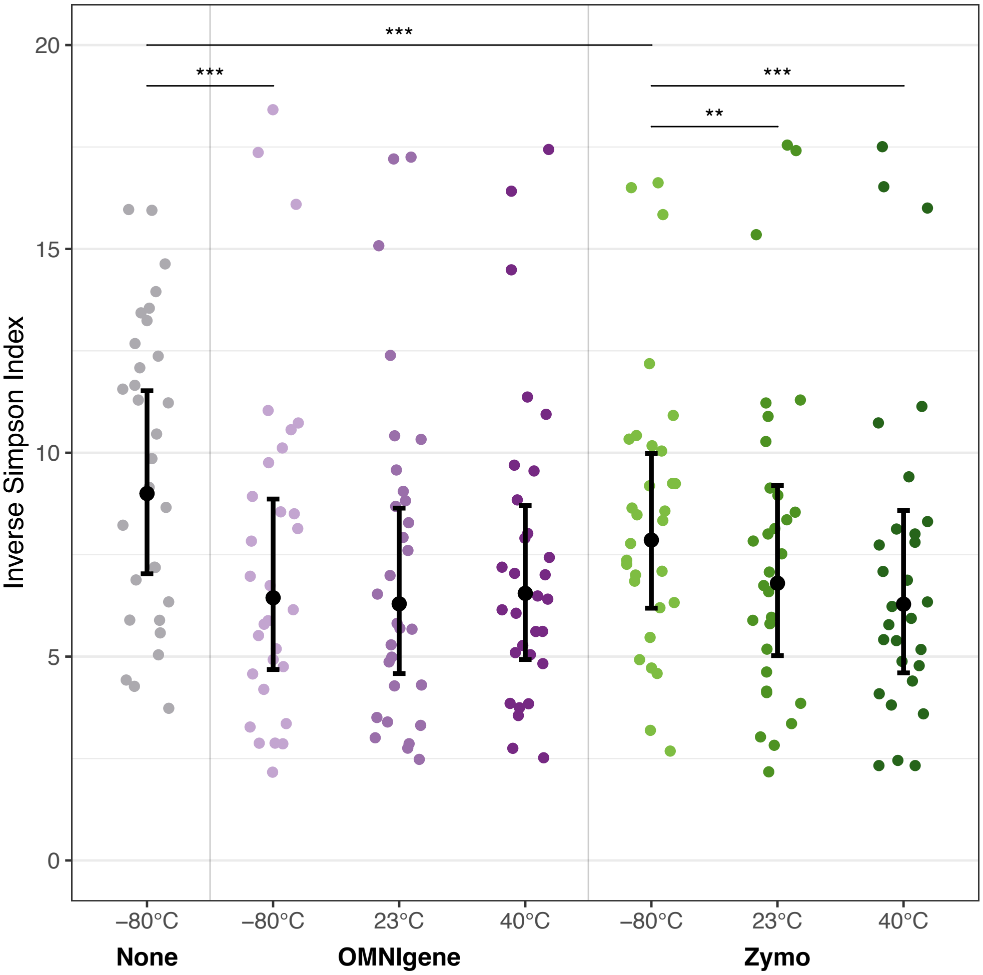


**Supplementary Figure 3: Metagenomic inverse Simpson index**

Genus-level metagenomic inverse Simpson index across samples from each condition. Scattered data points represent values from individual samples. Center points indicate estimated mean values from the GEE model. Whiskers indicate 95% upper and lower confidence intervals from the GEE model. *$p \leq 0.05$, **$p \leq 0.01$, ***$p \leq 0.001$.

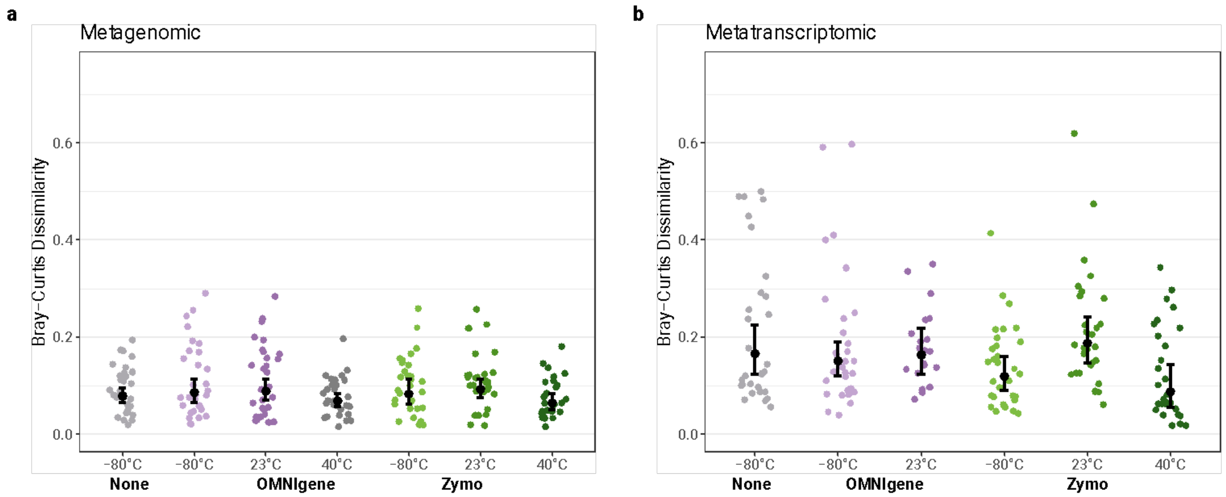


**Supplementary Figure 4: Within-condition metagenomic and metatranscriptomic Bray-Curtis dissimilarity**

Genus-level metagenomic (a) and metatranscriptomic (b) Bray-Curtis dissimilarity between technical replicates from each preservative condition. Each replicate from a given donor and condition was compared to the other replicates from the same donor and condition (for three total comparisons per donor and condition). There are no significant differences in Bray-Curtis within-condition dissimilarity. Scattered data points represent Bray-Curtis dissimilarity between pairs of samples. Center points indicate estimated mean values from the GEE model. Whiskers indicate 95% upper and lower confidence intervals from the GEE model.

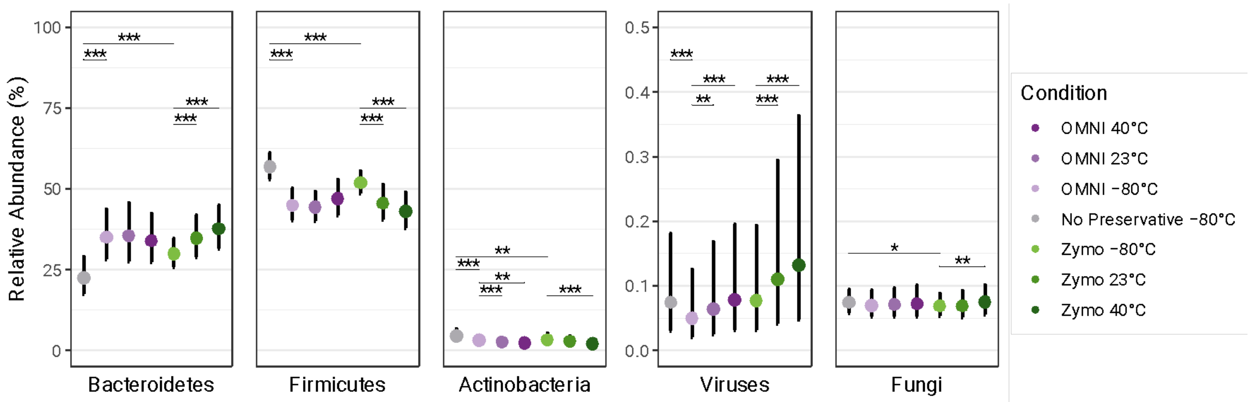


**Supplementary Figure 5: Metagenomic relative abundances of top bacterial phyla, viruses, and fungi**

Metagenomic relative abundance of the three most abundant bacterial phyla, viruses, and fungi across samples from each condition. Center points indicate estimated mean values from the GEE model. Whiskers indicate 95% upper and lower confidence intervals from the GEE model. *$p \leq 0.05$, **$p \leq 0.01$, ***$p \leq 0.001$.

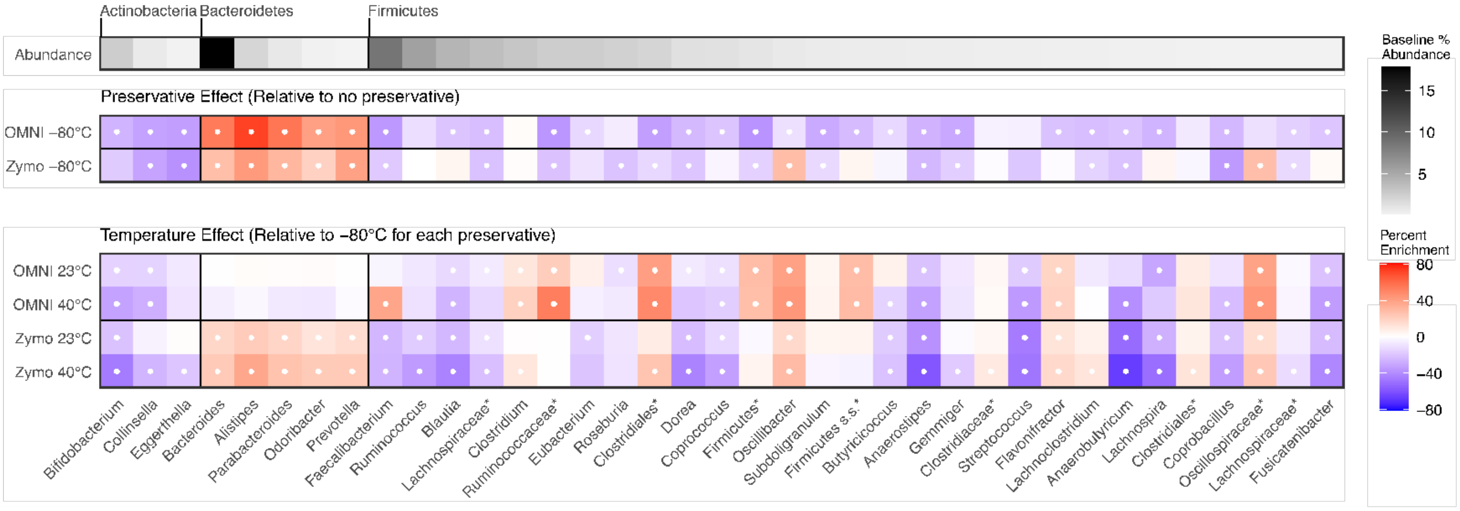


**Supplementary Figure 6: Metagenomic genus-level differential abundance across storage conditions**

Top bar indicates mean abundance of each genus in the no preservative, immediately frozen samples. Genera present at ≥ 0.1% modeled abundance in any condition are shown. Labels indicate phylum of each genus. Heatmap shows genus-level percent change in relative abundance between immediately frozen, preserved samples relative to immediately frozen, no preservative samples (middle rows) and change in relative abundance between samples exposed to heat in preservatives relative to immediately frozen, preserved samples (bottom rows). White circles indicate a significant percent change after Benjamini-Hochberg correction, FDR < 0.1. Asterisks in genus names indicate designated genera with no taxonomic name at the genus level (e.g. “miscellaneous Clostridiales”).

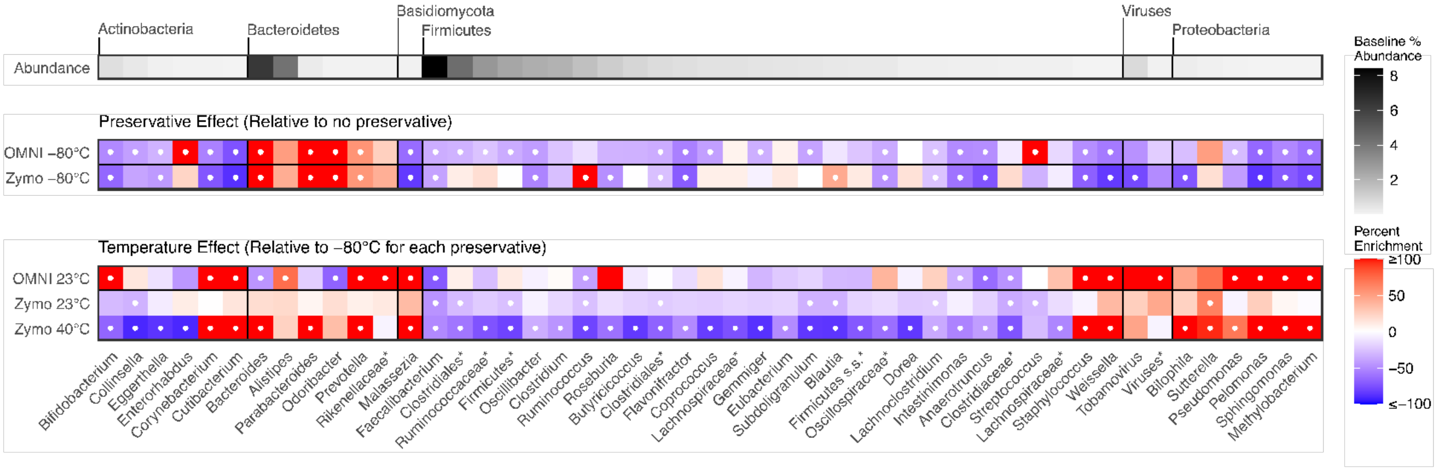


**Supplementary Figure 7: Metatranscriptomic genus-level differential abundance across storage conditions**

Top bar indicates mean abundance of each genus in the no preservative, immediately frozen samples. Genera present at ≥ 0.1% modeled abundance in any condition are shown. Labels indicate phylum of each genus. Heatmap shows genus-level percent change in relative abundance between immediately frozen, preserved samples relative to immediately frozen, no preservative samples (middle rows) and change in relative abundance between samples exposed to heat in preservatives relative to immediately frozen, preserved samples (bottom rows). White circles indicate a significant percent change after Benjamini-Hochberg correction, FDR < 0.1. Asterisks in genus names indicate designated genera with no taxonomic name at the genus level (e.g. “miscellaneous Clostridiales”).
